# Moderate increases in channel discharge are positively related to ecosystem respiration in forested Ozark streams

**DOI:** 10.1101/2021.01.12.426336

**Authors:** Allyn K. Dodd, Daniel D. Magoulick, Michelle A. Evans-White

## Abstract

The natural flow regime is considered the “master variable” in lotic systems, controlling structure and function at organismal, population, community, and ecosystem levels. We sought to estimate forested headwater stream metabolism across two dominant flow regimes (*Runoff* and *Groundwater*) in northern Arkansas and evaluate potential differences in, and drivers of, gross primary production, ecosystem respiration, and net ecosystem metabolism. Flow regimes differed in intermittency, substrate heterogeneity, hyporheic connectivity, and dominant water source (subsurface runoff vs. groundwater), which we expected to result in differences in primary production and respiration. Average daily gross primary production (GPP) and ecosystem respiration (ER) estimated from field data collected from May 2015-June 2016 tended to be greater in *Groundwater* streams. Respiration was positively related to discharge (R^2^= 0.98 p< 0.0001) and net metabolism became more heterotrophic with increasing average annual discharge across sites (R^2^= 0.94, p= 0.0008). Characterizing ecosystem-level responses to differences in flow can reveal mechanisms governing stream metabolism and, in turn, provide information regarding trophic state and energy inputs as efforts continue to determine global trends in aquatic carbon sources and fates.

## INTRODUCTION

The natural flow regime exerts primacy over water quality and quantity, habitat structure, disturbance regime, and, in turn, ecological processes and functions in lotic systems. Flow regime is characterized by the timing, duration, magnitude, frequency, and rate of change of water flowing through a channel over various temporal scales (Poff et al. 1997), arranging habitat space and thereby creating a unique template for life history strategies and community interactions (Southwood 1977, Poff and Ward 1990). Natural disturbances, such as flooding and drought, serve as life cycle prompts for many fishes and macroinvertebrates, whose reproductive cues are intimately linked with predictable, seasonal changes in flow (Poff and Ward 1989, Huryn and Wallace 2000, Humphries and Baldwin 2003, Lytle and Poff 2004). Flow regime is ultimately a byproduct of landscape-level processes and variation, as climate, topography, geology, vegetation, and soils interact to determine primary water sources (e.g. groundwater vs. subsurface runoff), quantity of within-channel flow, and geomorphology. Indeed, flow regime is typically a region- and land cover-specific phenomenon; streams reflect the diverse biomes that generate and sustain their flows as well as the relative contributions of groundwater, surface water, soil water, and precipitation (Hynes 1975, Poff et al. 1997, Carlisle et al. 2010).

Stream metabolism is an estimator of carbon (C) dynamics and an indicator of of nutrient cyclingand trophic status that is sensitive to natural and anthropogenic disturbances, revealing ecosystem-level responses to changes in hydrology and geomorphology. Metabolism is comprised of gross primary production (GPP) and ecosystem respiration (ER), which yields the net amount of carbon fixed into biomass, or net ecosystem metabolism (NEM) (Hall and Hotchkiss 2017). Metabolism can reveal whole-stream responses to landscape changes as well as predict potential bottom-up effects on higher trophic levels. Ecosystem metabolism is driven by proximal factors such as light and nutrients, which are influenced by the surrounding watershed (Bernot et al. 2010, Yates et al. 2013). The direct and indirect susceptibility of primary production and respiration to landscape-level variation makes it a useful metric for assessing impacts at the ecosystem level. Additionally, daily metabolism can vary temporally due to changes in light levels, organic matter inputs, algal biomass, and hydrology (Acuña et al. 2004, Roberts et al. 2007). Previous work assessing annual metabolism across multiple streams has focused primarily on the effects of biome and land use (Bott et al. 1985, Mulholland 2001, Bernot et al. 2010). The large dependence of other metabolism estimates on flow timing and magnitude (e.g. Uehlinger et al. 2003, Roberts et al. 2007, Qasem et al. 2018) suggests they will vary significantly across differing flow regimes within the same biome. Further, comparing function within and among hydrologic classifications provides insight into processes and variables controlling ecosystem function. A growing body of work has begun to address how stream metabolic regimes vary in time and space at multiple hierarchical scales (Appling et al. 2018, Koenig et al. 2019, Savoy et al. 2019), and a subset of these efforts have focused specifically on hydrologic influences impacting production and respiration (Jones et al. 1995, Dodds et al. 1996, Battin 1999, Uehlinger 2000, Uehlinger et al. 2003, Vilches and Giorgi 2010, Leggieri et al. 2013, Cook et al. 2015, Rovelli et al. 2017, Reisinger et al. 2017, Demars et al. 2019, O’Donnell and Hotchkiss 2019).

Existing conceptual models of headwater stream metabolism have posited that factors controlling metabolism differ by biome (Mulholland et al. 2001), land use category (Bernot et al. 2010), and season (Roberts et al. 2007). In reference systems, biome and season are considered the primary drivers of differences across streams (Bott et al. 1985, Mulholland et al. 2001, Hornbach et al. 2015). However, others have shown distinct hydroecological regions at hierarchical spatial scales characterized by significant variation in flow dynamics within a biome (Poff et al. 2006, Leasure et al. 2016). This variation arises from changes in geology and water sources across basins and sub-basins, which can result in differences in ecosystem function and carbon availability (Thoms and Parsons 2002).

Flow variability within a stream can be a determinant of annual metabolism, as flow extremes can exert a strong influence on organic matter movement through the system (Acuña et al. 2004, Roberts et al. 2007, Demars 2019). High flows can depress primary production (Uehlinger 2000, Uehlinger 2006) while increasing production rates in autumn by removing abscised leaves covering the benthos (Argerich et al. 2011). High flows can also influence respiration rates by reducing respiration initially due to loss of autotrophic biomass, then increasing rates as the autotrophic community recovers from scouring (Roberts et al. 2007, Izagirre et al. 2008). Consistently higher discharge, or higher discharge in one year compared to another, can depress primary production rates by preventing regrowth of algal biomass, while others have found clear relationships between hydrologic regime and benthic organic matter that supports respiration (Acuña et al. 2007, Demars 2019). Hot, dry summers that increase water temperature but reduce depth can support extensive algal production and may lead to an overall reduction in metabolic rates over summer in the absence of scouring floods (Izagirre et al. 2008).

Drying and flooding can both temporarily depress primary production and respiration, while the weeks following these disturbances are typically marked by high rates of production and respiration as algae recolonize the benthos (Uehlinger 2000, Uehlinger 2006). Specifically, the number of dry days, number of days experiencing high flows (defined as >75% average daily flow), and number of flood events affect production and respiration; the strength of this effect would be dependent upon the magnitude, frequency, and duration of the disturbance (Biggs et al. 2005, Palmer and Ruhi 2019).

Several natural flow categories have been identified for streams within the Ozark and Ouachita Interior Highlands in northern/western Arkansas, eastern Oklahoma, and southern Missouri (Leasure et al. 2016), but efforts to characterize these systems based on their unique hydrology in the field have only recently begun. Leasure et al. (2016) revealed distinct geographic areas demarcated by dominant flow types that are likely functionally unique. These flow classifications consist of subsurface runoff- and groundwater-fed systems, known as *Runoff Flashy* and *Groundwater Flashy* streams (hereafter *Runoff* and *Groundwater*), that are dominant in the Ozark and Boston Mountains ecoregions. Key differences between flow regimes are frequency and duration of low flow days, frequency of floods, channel substrate heterogeneity and size, and dominant water sources (e.g. groundwater versus subsurface runoff).

Both flow regimes experience flash floods marked by high magnitudes but short duration (Leasure et al. 2016). *Runoff* stream flow originates primarily from subsurface runoff originating from precipitation and tend to dry for several weeks during autumn. Mixing analysis has shown that, on average, 89 (± 6)% of *Runoff* base flows derive from precipitation in the form of interflow. *Groundwater* streams tend to flow perennially, with 79 (±3)% of base flow coming from groundwater contributions (Dodd et al. 2020) coming through the hyporheic zone. *Groundwater* streams tend to exhibit less variable annual flows (Leasure et al. 2016).

Additionally, channel geomorphology and dominant substrate are markedly different between flow regimes. *Groundwater* streams are dominated by a mixture of substrate types, dominated by sand, gravel, and pebble with some cobble and boulder mixed in; substantial bed movement is common during high flow events. Conversely, *Runoff* streambeds are often comprised of bedrock overlain in some areas by cobble and boulders. Substrate heterogeneity can alter near-bed flow and turbulence intensity, and in turn influence production rates (Cardinale et al. 2002), and a greater diversity of benthic substrate sizes may foster greater production in *Groundwater* streams. Our primary objective was to determine whether differences exist in stream gross primary production, ecosystem respiration, and net ecosystem production between *Runoff* and *Groundwater* streams and characterize natural variation in ecological-flow responses across seasons. We expected average GPP to be greater in *Groundwater* streams, as these streams tend to exhibit perennial flow, have relatively stable hydrology over the year, and exhibit low turbidity. We expected ER to be greater in *Groundwater* systems as well given that the heterogeneous benthic substrate may result in higher rates of respiration by providing spaces for organic matter to settle and pockets in the benthic and hyporheic zones for heterotrophic microbes to colonize.

We also sought to confirm that differences exist in stream discharge and flow metrics. We hypothesized that *Groundwater* streams would have greater annual discharge since these streams do not dry and have more sustained, less variable flows from groundwater.

We also examined potential drivers of metabolism across sites. We predicted positive relationships between GPP and light, nutrients (total N and total P), algal biomass, and a negative relationship between GPP and discharge, number of high flow days, number of floods, number of low flow days, and number of no flow days. We hypothesized that positive relationships would exist between ER and biofilm ash-free dry mass, nutrients, floods, high flow days, and discharge (Figure 1).

**Figure 1.**
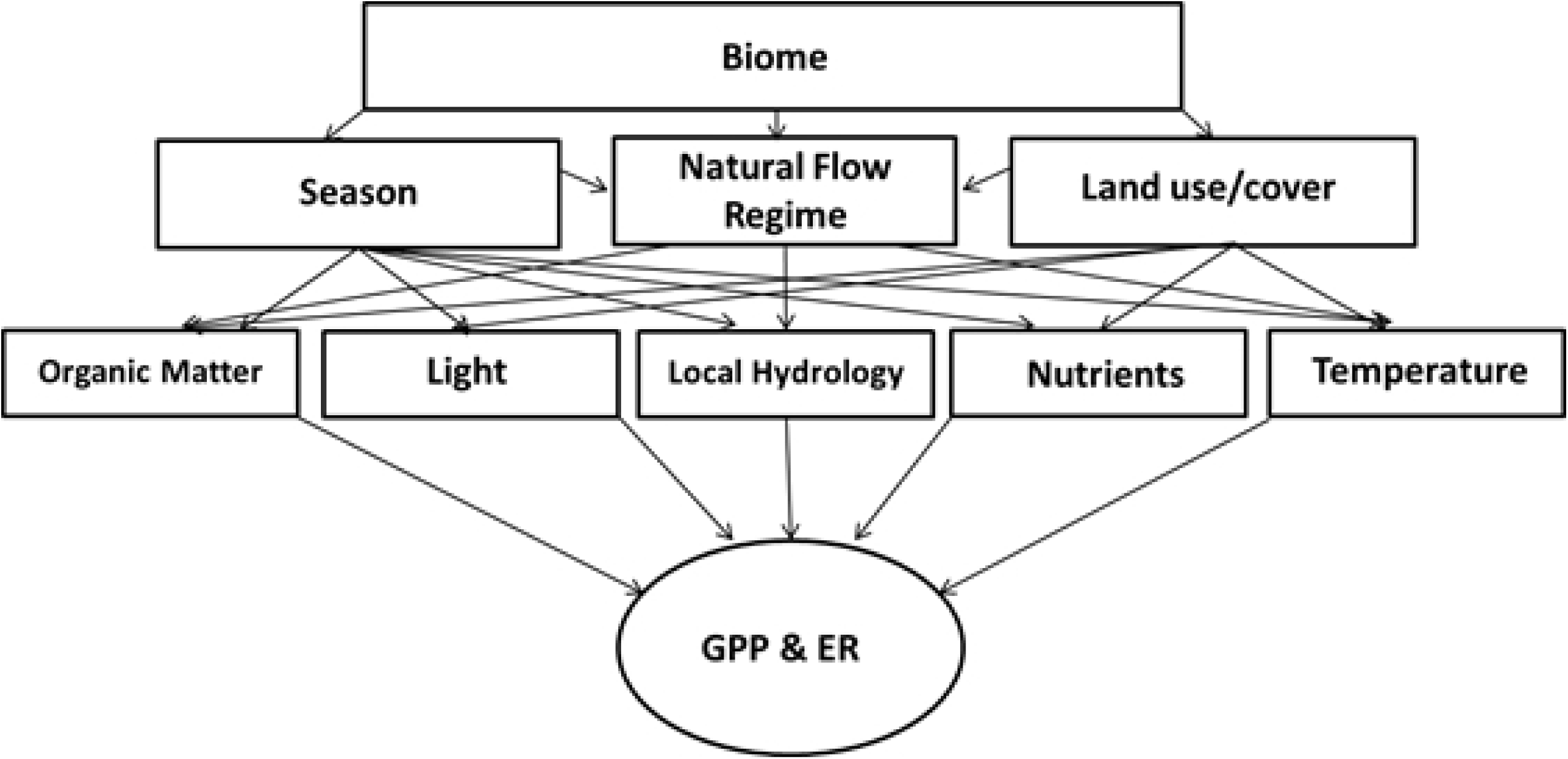
Conceptual diagram illustrating factors controlling stream metabolism, which incorporates the natural flow regime of an area as influenced by biome, season, and land use. This, in turn, can influence hydrologic responses to land use change. GPP and ER denote gross primary production and ecosystem respiration, respectively.

## METHODS

### STUDY SITES

This study was conducted in six temperate, minimally impacted headwater streams in the deciduous forests of Arkansas (Figure 2). We chose three *Groundwater* streams and three *Runoff* streams categorized based on flow classifications modeled by Leasure et al. (2016). Study sites were chosen based on membership in *Runoff Flashy* or *Groundwater Flashy* flow classes as well as stream surface area, surrounding land cover, and presence of downstream USGS stream gauges. Streams selected for the study were all headwater systems with forested land cover ranging from 84 to 95% of total watershed area (Stroud Water Research Center 2017) (Table 1).

**Table 1.**
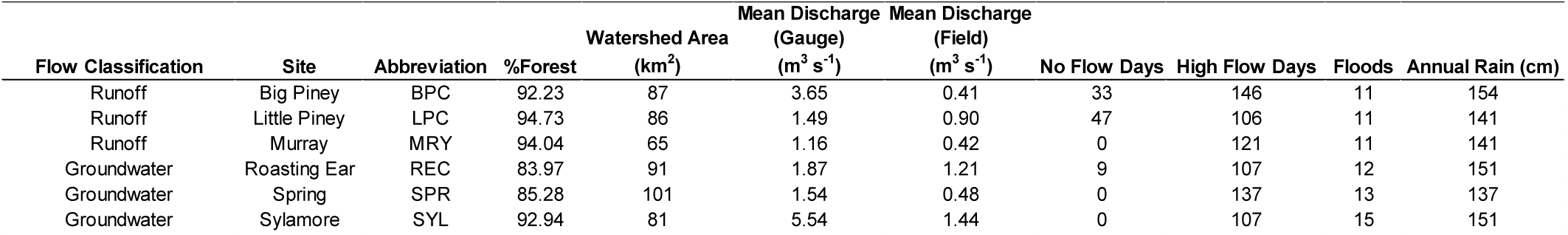
Site abbreviations and flow metrics for May 2015 through June 2016. Discharge estimates reflect year-round measurements calculated from daily gauge data (Gauge) or from discharge measured monthly in the field, which were taken at or near base flow (Field).

**Figure 2.**
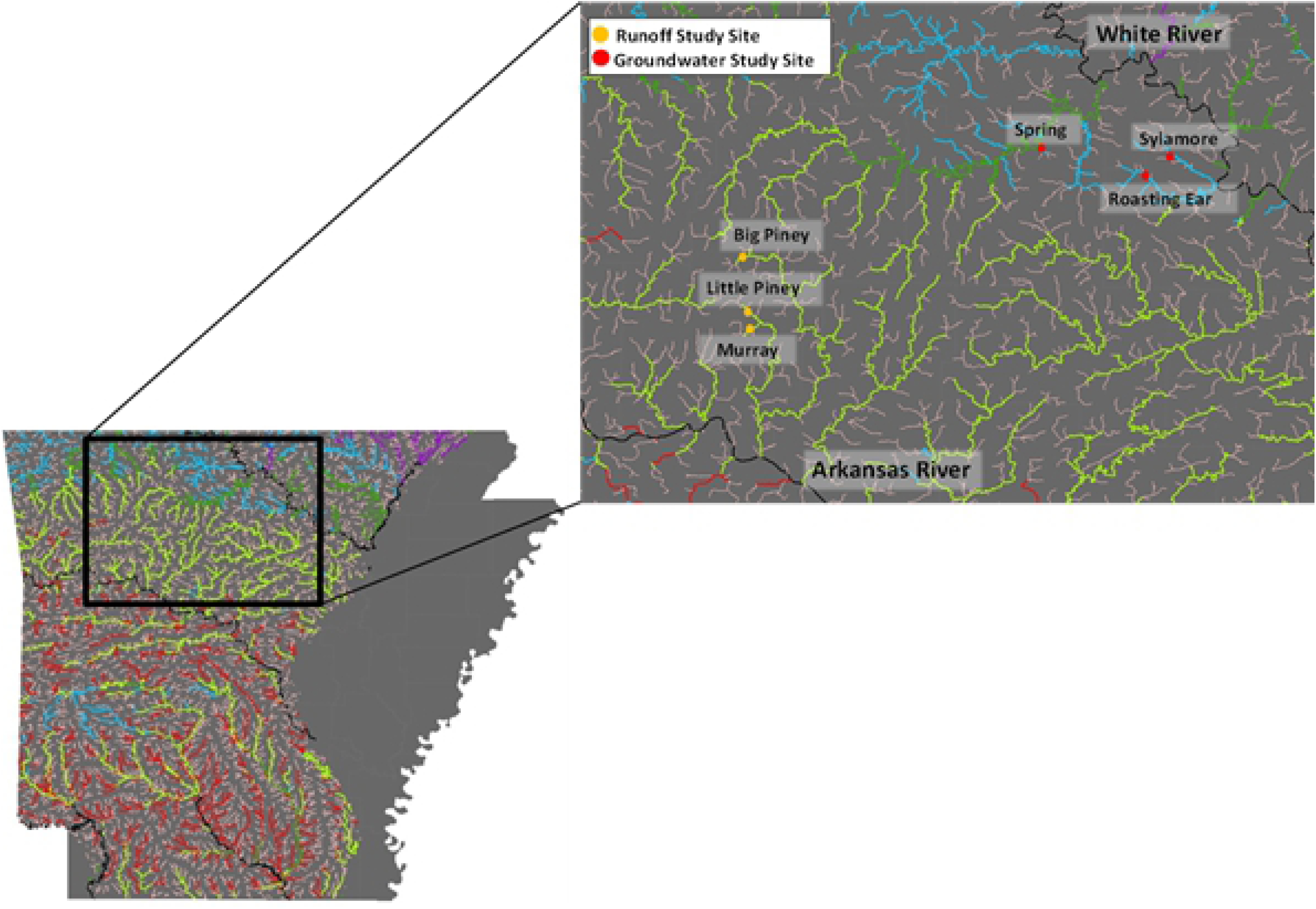
Map of flow regimes in the Ozark and Ouachita Interior Highlands based on Leasure et al. (2016). Highlighted area shows individual study sites sampled from 2015-2016 across northern Arkansas. Teal lines represent *Groundwater* streams. Light green lines irepresent *Runoff* streams.

All streams are located within the Ozark National Forest in northern Arkansas and were chosen based on the amount of surrounding forest in the watershed and a lack of tributaries or ephemeral drains feeding into accessible reaches to prevent extraneous variables influencing GPP and ER. Vegetation surrounding both stream types was primarily oak and hickory trees forest (Woods et al. 2004). While all streams had similar surrounding landcover and vegetation, the geology underlying *Runoff* and *Groundwater* streams differ. *Groundwater* streams are found within a highly dissected limestone plateau. Conversely, the *Runoff* streams are found in an area dominated by green/gray shale and sandstone. Vegetation surrounding both stream types was primarily oak and hickory trees forest (Woods et al. 2004, Chapman et al. 2006, Stephenson et al. 2007).

### METABOLISM RATES

We calculated reach-scale metabolism using the open-channel single-station method (Odum 1956, Riley and Dodds 2012). Dissolved oxygen (DO) and temperature were measured every 15 minutes by Hydrolab DS5X multiparameter sondes (Hach Company, Loveland, CO) from May 2015 to June 2016 in a well-mixed area at the bottom of each study reach. Data were corrected as necessary by comparing with DO concentrations determined via Winkler titrations (Dodds et al. 2018). Reaeration coefficients as estimates of air-water gas exchange were determined via propane release in five out of six streams, while nighttime regression was utilized in one *Runoff* stream, Murray Creek (Hall and Hotchkiss 2017). Propane release was necessary in five streams because nighttime regressions yielded significant relationships between ER and air-water gas exchange, or reaeration, coefficients (K_600_), whereas no such relationship was present at Murray Creek. Corrections for groundwater contributions to reaches receiving appreciable inputs were made according to Hall and Tank (2005). Photosynthetically-active radiation (PAR) measurements were logged concurrently with metabolism parameters using an Odyssey light meter positioned in an area near the stream with open canopy. Stream metabolism was estimated based on diel changes in DO, temperature, depth, and light measurements; we used R package StreamMetabolizer (Appling et al. 2018) to solve for GPP and ER utilizing a single-station metabolism maximum likelihood model with the core equation:

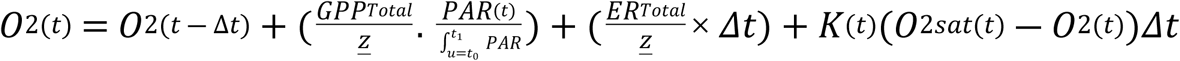

where t is time and Δt is the time step between measurements (15 minutes), *<u>z</u>* is mean reach depth, 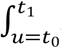 *PAR* is daily photosynthetically-active radiation, and K_(t)_ is air-water gas exchange corrected for temperature. Estimates of GPP_Total_ and ER_Total_ fitted by StreamMetabolizer yielded daily rates for every day that a sonde was deployed at each stream (from 158 to 215 days). Data were not collected every day of the 422-day study due to flash floods and/or unwadeable conditions, drying, and/or equipment failure. Seasonal GPP and ER were calculated by averaging daily rates from the beginning of the respective season (e.g. equinox or solstice). Overall average daily rates were calculated by averaging daily rates over the study period. Summing daily rates by season would hinder comparisons since we did not have equal numbers of sampling dates across streams, so averages of daily rates over each season are reported. Average daily rates for each flow class were computed by computing the mean of the three streams in that flow class’ overall average daily rates.

### PHYSICOCHEMICAL VARIABLES

To evaluate relationships between metabolism and flow metrics, we quantified high flow days, number of floods, and number of days with no flow. High flow days were defined as exceeding the 75^th^ percentile of mean annual discharge at a site. Floods were quantified as distinct hydropeaks greater than 100% of mean annual flow calculated from gauge data and upstream-downstream discharge relationshipsbetween USGS gauges and discharge measured in each study reach. We measured discharge in each reach monthly from May 2015 to June 2016 using velocity-area gauging (Gore 2006). We measured median diameter (d50) of randomly selected stones across each discharge transect. One stone was randomly selected at each meter across every discharge transect during four spring and summer 2016 monthly sampling events.

Persulfate digests of unfiltered water samples taken at base or near-base flow during monthly sampling events were followed by colorimetric analyses to determine nutrient concentrations. Total nitrogen (TN) was measured monthly by automated cadmium reduction on a Lachat Quikchem 8500 (Hach Company, Loveland, CO). Total phosphorus (TP) was measured monthly using the ascorbic acid method (APHA 2005).

Canopy cover was determined for each stream channel once in summer and once following abscission using a densiometer to calculate percent coverage.

### PERIPHYTON

For algal biomass, we collected six cobbles per reach at six equidistant transects down the stream reach. Monthly algal biomass was determined beginning in summer 2015 by scrubbing rocks of algae byextracting chorophyll *a* each from algal slurry in ethanol, then wrapping with aluminum foil to determine rock area from mass-area relationships (Steinman et al. 2006).

### STATISTICS

We used 2-way repeated-measures ANOVA to determine whether differences existed in metabolism, flow metrics, and periphyton biomass and biofilm ash-free dry mass between flow regimes by season. Linear regressions were employed to examine relationships between daily metabolism variables (GPP, ER, and NEP), physicochemical parameters (e.g. total nitrogen and phosphorus, temperature), biological metrics (e.g. chlorophyll *a*, biofilm ash-free dry mass), and flow metrics (e.g. gauge-calculated discharge, discharge measured by hand in each reach, number of low flow days, number of high flow days, coefficient of variation of daily flow, number of floods). We evaluated relationships between variables with both gauge-calculated discharge, which yielded estimates of flow for all days of the study including extreme floods and times during which sondes were not deployed, and discharge measured in the reach monthly, which represented conditions under which sondes were deployed and did not include extreme high flows (Table 1). All statistical analyses were performed in R version 3.4.3. Statistical significance threshold was p ≤ 0.05.

## RESULTS

### METABOLISM

There was no significant effect of flow regime on average daily GPP across seasons (Table 2), though *Groundwater* streams tended to show greater GPP all year except during winter (Figure 3). Average daily GPP was highest in summer 2016 in *Groundwater* streams, while *Runoff* streams exhibited the greatest average daily GPP in summer 2015. Total GPP ranged from 99.7 to 435.1 g O_2_ m^-2^ y^-1^ in *Runoff* streams (days of record= 158-188 days), and 289.5 to 405.4 g O_2_ m^-2^ y^-1^ in *Groundwater* streams (days of record= 159-202 days). Big Piney and Murray Creek, two *Runoff* streams, had fairly constant rates of GPP over the study year, with very small increases in GPP during the spring and summer. Conversely, the third *Runoff* site, Little Piney, showed marked increases in production during the spring and summer months, with daily production remaining high until the stream dried in October. Similarly high daily production rates were found at Sylamore, a *Groundwater* stream, throughout the spring and summer, though production decreased from July to September. The other two *Groundwater* sites exhibited spring and summer production rates like Big Piney and Murray, both *Runoff* streams. However, *Groundwater* stream production appeared to be stimulated by floods that occurred at the end of 2015, while *Runoff* streams showed little to no such response. Additionally, all *Groundwater* streams and one *Runoff* (Little Piney) revealed greater primary production in summer 2016 compared to the previous year.

**Table 2.** Results of repeated measures Analysis of Variance (ANOVA) for variables of interest. * denote statistical significance.

| Dependent Variable<br>(2-way RM-ANOVA) | F-statistic | p- value |
| --- | --- | --- |
| **GPP** |  |  |
| Flow | $F_{(1,12)} = 1.06$ | 0.32 |
| Season | $F_{(3,12)} = 0.96$ | 0.31 |
| Flow x Season | $F_{(5,12)} = 1.26$ | 0.48 |
| **ER** |  |  |
| Flow | $F_{(1,12)} = 0.08$ | 0.79 |
| Season | $F_{(3,12)} = 0.18$ | 0.91 |
| Flow x Season | $F_{(5,12)} = 1.05$ | 0.39 |
| **Discharge** |  |  |
| Flow | $F_{(1,12)} = 2.15$ | 0.17 |
| Season | $F_{(3,12)} = 2.86$ | 0.04* |
| Flow x Season | $F_{(5,12)} = 1.41$ | 0.29 |
| **Total Nitrogen** |  |  |
| Flow | $F_{(1,12)} = 0.12$ | 0.75 |
| Season | $F_{(3,12)} = 2.35$ | 0.17 |
| Flow x Season | $F_{(5,12)} = 2.54$ | 0.21 |
| **Total Phosphorus** |  |  |
| Flow | $F_{(1,12)} = 0.04$ | 0.85 |
| Season | $F_{(3,12)} = 2.35$ | 0.17 |
| Flow x Season | $F_{(5,12)} = 0.30$ | 0.61 |
| **Algal Biomass** |  |  |
| Flow | $F_{(1,12)} = 1.39$ | 0.27 |
| Season | $F_{(3,12)} = 8.34$ | 0.0004* |
| Flow x Season | $F_{(5,12)} = 2.78$ | 0.09 |
| **Ash-Free Dry Mass** |  |  |
| Flow | $F_{(1,12)} = 18.56$ | 0.0005* |
| Season | $F_{(3,12)} = 6.60$ | 0.004* |
| Flow x Season | $F_{(5,12)} = 2.21$ | 0.13 |

**Figure 3.**
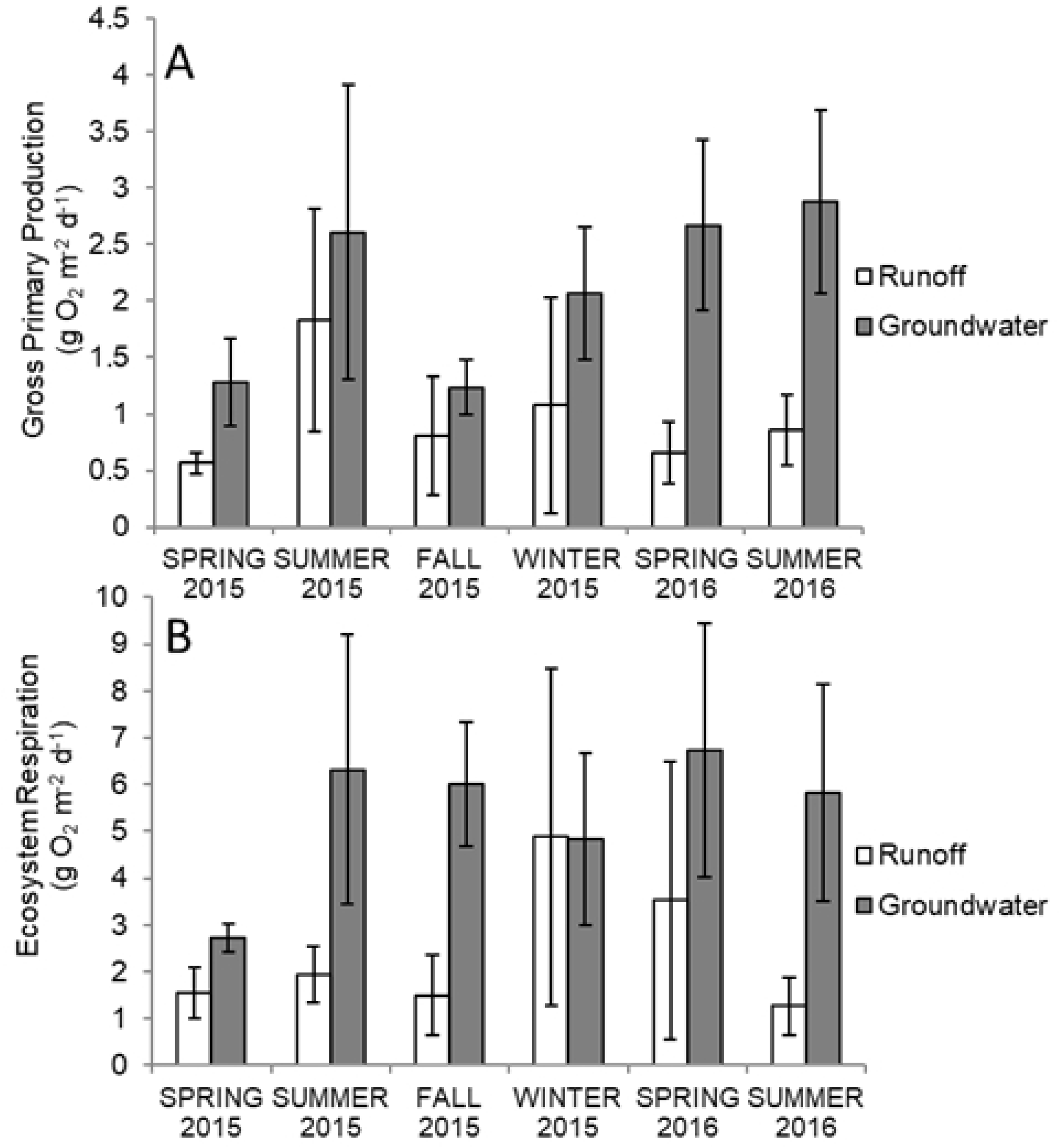
Seasonal average daily gross primary production (A) and respiration (B) in *Runoff* (white) and *Groundwater* (gray) streams. Error bars denote ± 1 standard error. n= 3 per flow regime.

We found no difference in average daily ER between flow regimes across seasons, though ER also tended to be greater in *Groundwater* streams. Average daily ER was greatest in spring 2016 across streams. Average daily GPP in *Runoff* streams was 1.2 (± 0.54) g O_2_ m^-2^ d^-1^, while *Groundwater* streams averaged 1.9 (± 0.27) g O_2_ m^-2^ d^-1^. Daily ER in *Runoff* streams averaged −2.1 (± 0.99) g O_2_ m^-2^ d^-1^ and −5.2 (± 1.0) g O_2_ m^-2^ d^-1^ in *Groundwater* streams. Daily rates of GPP and ER for each stream are shown in Figure 4. Total ER in *Runoff* streams ranged from −151.4 to −771.2 g O_2_ m^-2^ y^-1^ and from −280.2 to −1340.3 g O_2_ m^-2^ y^-1^ in *Groundwater* streams. Peaks in respiration rates over the year varied across all six sites. While Big Piney and Murray (*Runoff* streams) had low respiration rates (similar to GPP), trends in respiration in Little Piney were similar to those observed in *Groundwater* streams. Respiration was especially high in Little Piney and *Groundwater* streams during the summer of 2015 and spring 2016, though *Groundwater* streams overall showed greater rates over the year and elevated respiration extended well into the autumn of 2015. Respiration was also stimulated by the heavy flooding that occurred in December 2015. In *Groundwater* streams, respiration was greater in summer 2016 than the previous year, while respiration-though low overall in comparison-was greater in summer 2015 than 2016 in *Runoff* streams.

**Figure 4.**
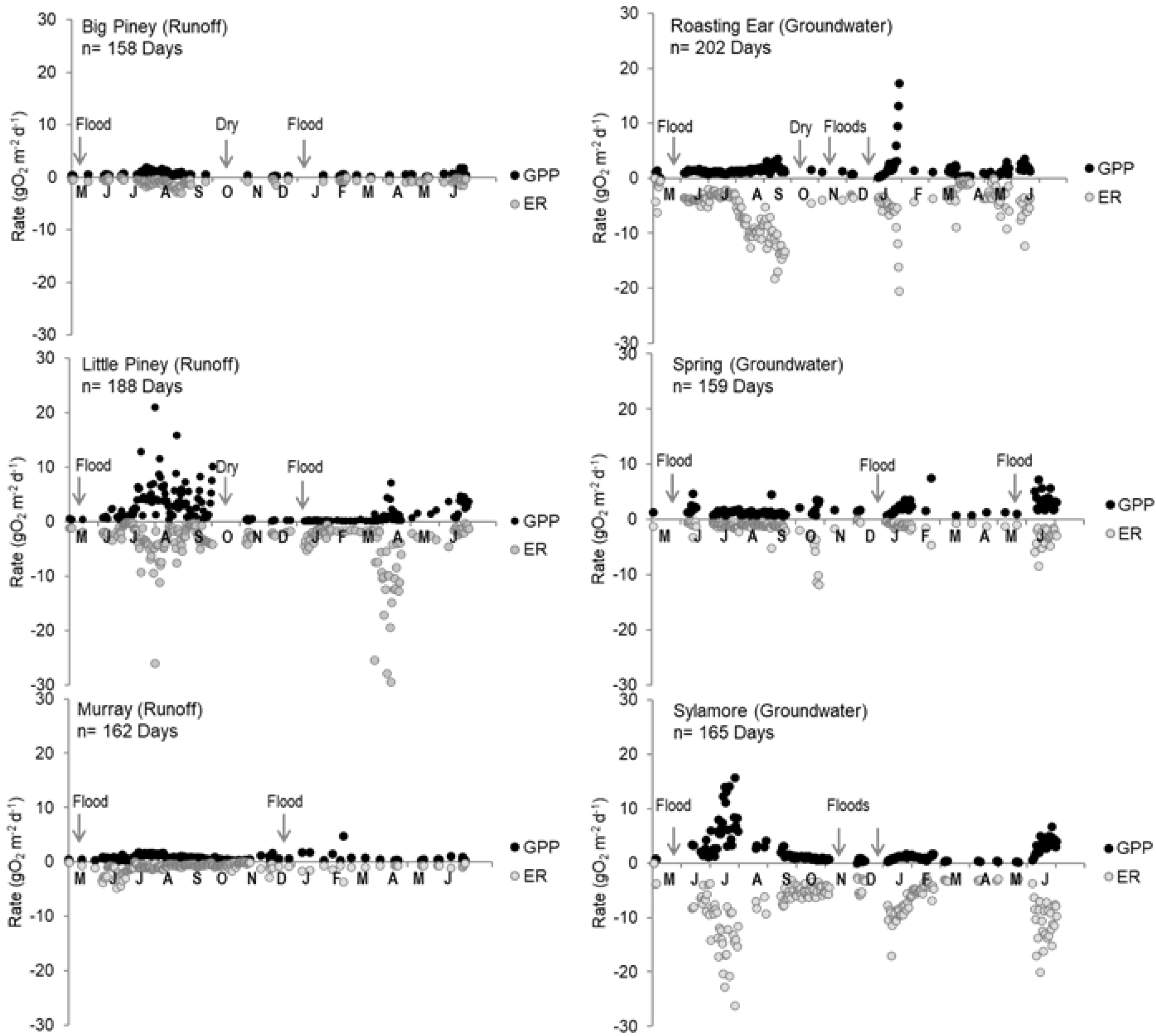
Daily rates of gross primary production (GPP) (black) and ecosystem respiration (ER) (gray) in Big Piney, Little Piney, Murray, Roasting Ear, Spring, and Sylamore from May 2015 to June 2016. *Runoff* streams are shown in panels on the left, *Groundwater* streams are represented in panels on the right. n= number of days of record at each site. “Flood” denotes large floods that prevented data collection.

### PHYSICOCHEMICAL VARIABLES

Stream discharge did not differ significantly across flow regimes but differed by season (F_(5, 12)_= 2.86, p= 0.04) (Table 2), with discharge peaking in the spring following substantial heavy winter rains that resulted in extreme flash floods in four out of six sites (Figure 5).

**Figure 5.**
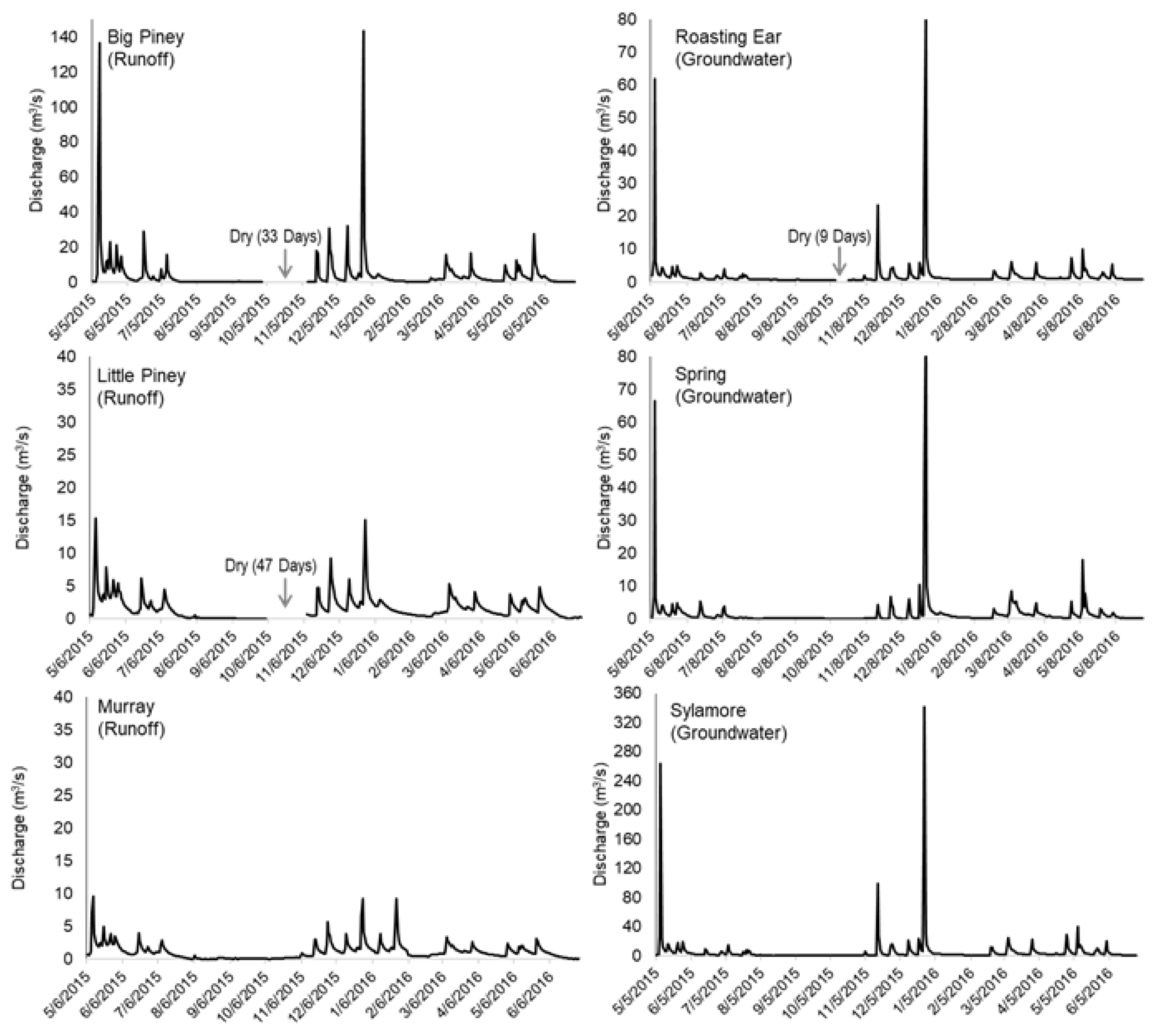
Hydrographs for study sites from May 2015 to June 2016. Note differences in y-axis scales; scales were not standardized to preserve details of individual site hydrology.

Discharge tended to be greater in *Groundwater* streams over the study period; however, these systems showed substantially elevated flows compared to *Runoff* streams during the spring across both 2015 and 2016. Between March and June 2016, *Groundwater* streams experienced one to two more high-flow events than *Runoff* streams. *Runoff* streams experienced two moderate storm events during summer 2015, while *Groundwater* streams experienced three smaller events during the same period. Little to no rain fell across Arkansas from early August to November 7, 2015, causing two *Runoff* streams and one *Groundwater* stream to dry. However, *Runoff* streams that dried experienced longer drought periods, as Little Piney and Big Piney dried for 47 and 33 days, respectively, whereas Roasting Ear (a *Groundwater* stream) dried for only 9 days. Precipitation inputs did not differ significantly across flow regimes (p= 0.85). Over the year, *Runoff* sites received 145 ± 4 cm of rainfall, while *Groundwater* sites received 146 ± 4 cm. Flow metrics at each site are listed in Table 1.

Greater discharge stimulated respiration (R^2^= 0.98, p< 0.0001) and drove streams to be more heterotrophic over the year (R^2^= 0.94, p= 0.0008) (Figure 6). Discharge is often related to watershed area and stream size, though we did not find a relationship between watershed area and discharge (R^2^= 0.004, p= 0.89). Cooler *Groundwater* streams had greater rates of primary production; GPP was negatively related to water temperature across sites (R^2=^ 0.63, p= 0.04). We found no further relationships between GPP or ER and physicochemical or biological variables.

**Figure 6.**
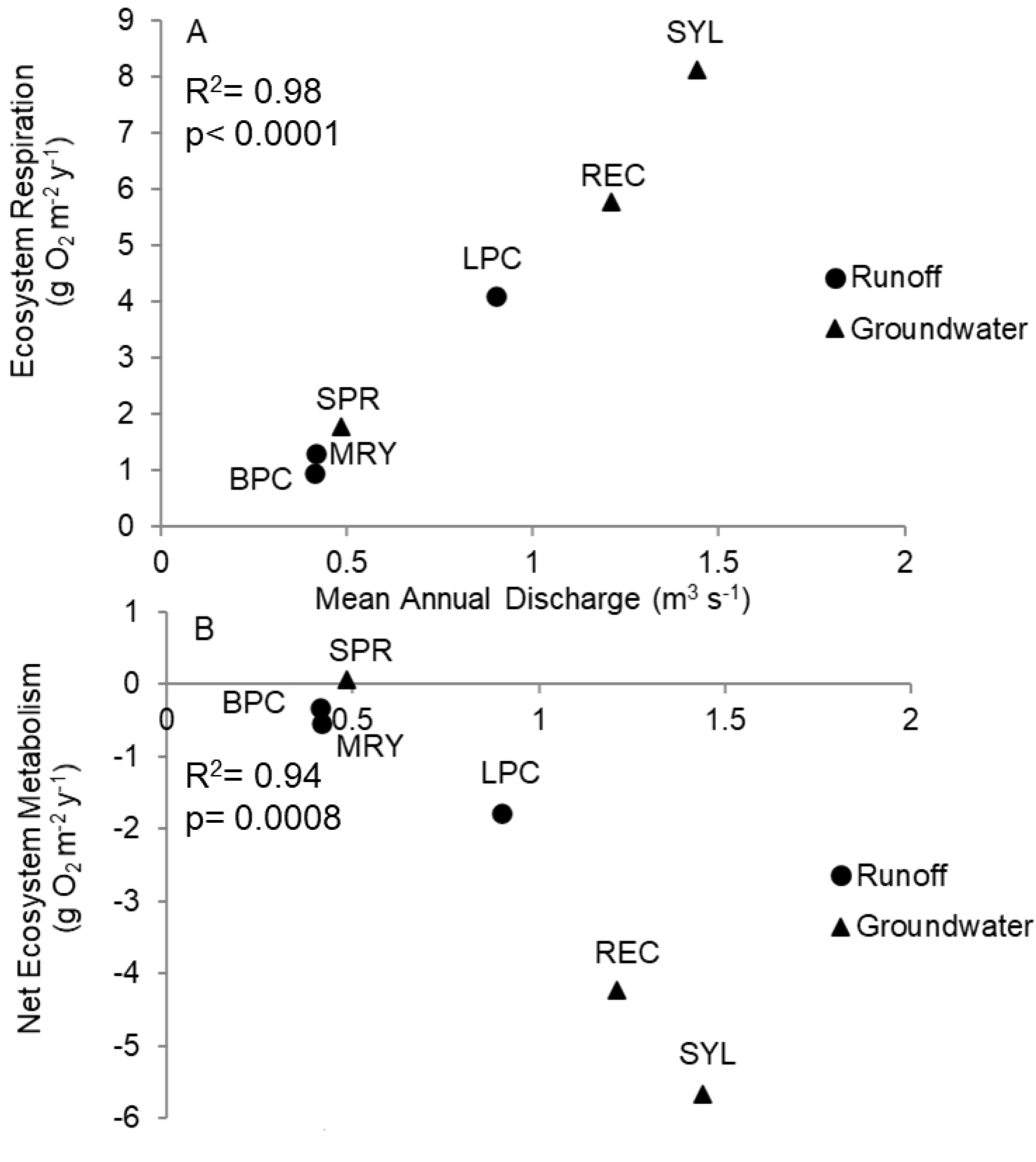
Ecosystem respiration (A) and net ecosystem metabolism (B) compared with mean annual discharge measured in the field across flow regimes. These discharge measurements were taken only when streams were wadeable and thus represent variation in discharge across excluding high flows.

*Runoff* stream substrate types were primarily bedrock and cobble, while *Groundwater* streams were dominated by pebbles with a variety of other substrate sizes present (i.e. sand, gravel, cobble, and some boulders). Median particle size (d50) was 19 <u>mm</u> (range= 10-82 mm) in *Runoff* streams and 13 mm (range= <1-110 mm) in *Groundwater* streams.

Nutrient concentrations were not significantly different across sites and seasons, though both TN and TP tended to be greater in *Groundwater* streams (Table 2). Total N averaged 0.10 ± 0.03 mg/L in *Runoff* streams and 0.56 ± 0.26 mg/L in *Groundwater* streams. Mean total P was 6.21 ± 0.63 μg/L in *Runoff* streams and 8.70 ± 1.38 μg/L in *Groundwater* streams.

We explored potential relationships between watershed land cover and nutrient concentrations to determine whether even forested systems could be experiencing impacts of surrounding anthropogenic activity. Instream phosphorus concentrations decreased as surrounding forested land cover increased within a watershed (R^2^= 0.77, p= 0.01). We observed a similar negative trend with respect to TN concentrations and forested land cover, though this relationship was not statistically significant (R^2^= 0.56, p= 0.09).

### PERIPHYTON AND ORGANIC MATTER

Algal biomass was not significantly different between flow regimes, though there was a seasonal effect (F_(5, 12)_= 8.33, p= 0.0004) (Table 2) and algal biomass tended to be greater in *Groundwater* streams throughout the year. Algal biomass peaked in the spring of 2016 in all streams. Groundwater stream algal biomass over the year was 3.4 ± 1.4 *µ*g/cm^2^ and Runoff stream algal biomass was less than half of that, averaging 1.5 ± 0.1 µg/cm2. Organic matter measured as ash-free dry mass differed across seasons (F_(3, 12)_= 6.60, p= 0.004) and was greater in *Groundwater* streams across all seasons except winter (F_(1,12)_= 18.56, p= 0.0005) (Figure 7). Similar to algal biomass, mean *Groundwater* stream biofilm ash-free dry mass was 11.7 ± 1.5 mg, while Runoff streams yielded 5.9 + 0.5 mg.

**Figure 7.**
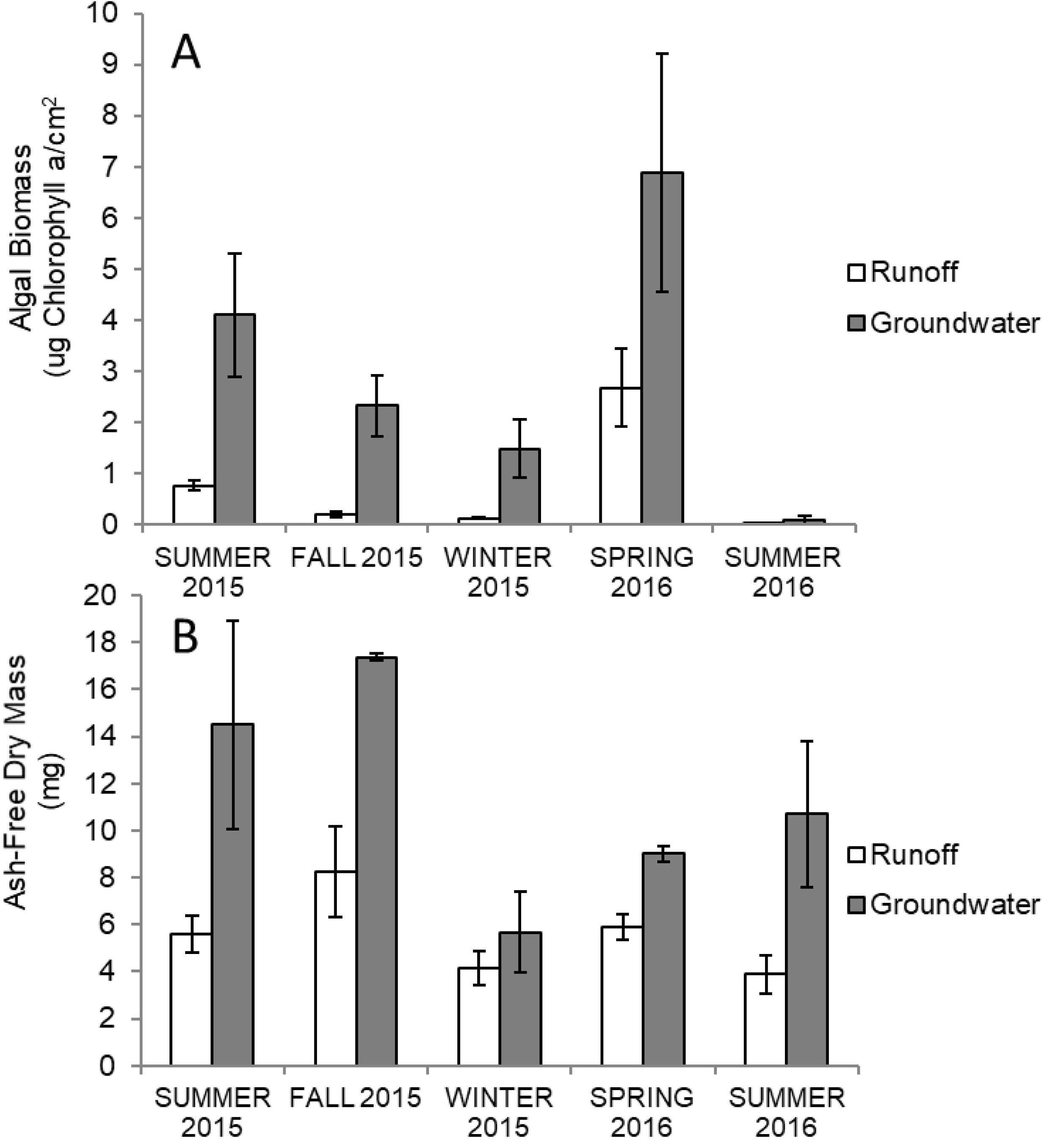
Seasonal algal biomass (A) and biofilm slurry ash-free dry mass (B) in *Runoff* and *Groundwater* streams. Error bars denote ± 1 standard error. n= 3 per flow regime.

We evaluated whether instream nutrient concentrations could be driving differences between algal biomass and ash-free dry mass. Neither TN nor TP were related to algal biomass (TP: R^2^= 0.47, p= 0.13, TN: R^2^= 0.38, p= 0.19) or ash-free dry mass (TP: R^2^= 0.10, p= 0.56, TN: R^2^= 0.39, p= 0.18).

## DISCUSSION

### METABOLISM

Primary production tended to be greater in *Groundwater* streams during spring and summer 2016, possibly resulting from *Groundwater* streams’ greater spring flows, slightly smaller, more mobile benthic substrate, and limestone karst rather than shale and sandstone that characterize the geology of *Runoff* sites. Discharge was greater in *Groundwater* streams during the spring, resulting from heavy rains that fell at the end of 2015. *Groundwater* streams experienced floods of much greater magnitude, and experienced elevated flows for the duration of the spring that were not the product of differences in rainfall between *Runoff* and *Groundwater* sites. While floods can depress primary production through scouring of the benthos (Grimm and Fisher 1984, Uehlinger et al. 2003, Roberts et al. 2007), moderately elevated discharge can stimulate production by reducing competition for light and nutrients through continuous thinning of the biofilm while more nutrients are transported downstream (Stevenson 1990, Humphrey and Stevenson 1992). Further, channel substrate can influence GPP; in particular, a greater diversity of substrate sizes, as found in *Groundwater* streams, has been shown to increase primary production (Cardinale et al. 2002). The highly variable rock sizes included more mobile substrate that may have further stimulated GPP by providing more surface area for algal colonization and allowing for slight bed movement during spring rain events, which contrasts with the relatively homogeneous boulder and bedrock substrate of *Runoff* streams. Greater GPP in *Groundwater* streams may be a byproduct of channel substrate coupled with greater water clarity during the spring’s higher flows; the benthos of each *Runoff* stream was difficult to observe if the water was more than approximately 0.5 meters deep (such as during the spring) due to the presence of minerals from glauconitic, easily-weathered green shale common to the Boston Mountains ecoregion (Caplan 1957, Caplan 1960), where *Runoff* streams are the dominant flow type. Conversely, *Groundwater* streams were clear year-round. While *Runoff* streams were warmer than *Groundwater* streams and contained more stable substrate for algal colonization, mineral effects on water clarity may have restricted primary production potential.

Similar to patterns reported in other forested systems and larger rivers, daily *Groundwater* stream GPP was greatest in spring and summer 2016, while summer 2015 had the greatest average GPP for *Runoff* streams (Mulholland et al. 2001, Acuña et al. 2004, Genzoli and Hall 2016). However, these trends differ from Roberts et al. (2007) in that GPP at Walker Branch was greatest during the spring and comparatively low during two consecutive summers. Uehlinger (2006) also reported GPP to be greatest in May over a 15-year period and lower in summers. *Runoff* stream GPP was lowest in spring 2015, coinciding with maximum algal biomass that may have reduced production rates due to competition. Daily *Groundwater* stream GPP was lowest in autumn 2015, which may have arisen from competition between benthic producers and microbes for resources, as reduced flows are common during early-to-mid autumn in Arkansas. It is worth noting that maximum algal biomass and production rates will not necessarily coincide. Algal biomass represents the state/structure of the algal community at a specific timepoint, and structure does not always directly reflect function (such as rates of production). At high levels of algal biomass, competition for light and nutrients may slow production rates (Sumner and Fisher 1979, Morin et al. 1999). While algae are allocating energy to maximize resource exploitation or resist the deleterious impacts of nutrient or light limitation due to lower competitive ability, there is less energy available for production, thus reducing primary production despite high biomass (McCormick 1996).

Respiration tended to be greater in *Groundwater* streams all year except during winter. This trend may have resulted from increased microbial activity post-abscission during the fall and a lack of large flood events through the summers and autumn. *Runoff* streams dried around the time of abscission, but even when rainfall replenished channel flow, respiration rates remained lower; a lack of substantial benthic storage in *Runoff* streams may have led to a net export of organic matter once rains returned in November. More variable substrate size and a hyporheic corridor in *Groundwater* streams could have facilitated greater community respiration rates by providing spaces for organic matter storage and colonization of heterotrophic microbes (Demars 2019).

While our analyses did not reveal statistically significant differences in metabolism across flow regimes, *Groundwater* streams tended to have greater rates of GPP and ER that are likely biologically relevant to instream food webs and carbon dynamics. Production rates were two times greater in *Groundwater* streams in nearly every season, while respiration was, in some seasons, three and four times greater. *Groundwater* streams held significantly more biofilm ash-free dry mass, revealing that even marginally greater production and respiration rates can have noticeable effects on basal resources and the biological community. The tendency for *Groundwater* streams to have higher rates of GPP and ER through much of the year almost certainly influences algal and consumer community structure. Others have found relationships between metabolism and fish assemblages (Munn et al. 2020), and further work measuring invertebrate and fish assemblages across systems simultaneously with metabolism would provide insight into the potential link between biologically significant differences in stream metabolism and food web effects. Thus, while our statistical results did not detect an effect of flow regime on ecosystem function, that does not discount flow regime as an important variable shaping stream metabolic regime.

Our results show a stimulatory effect of discharge on ER, just as others have found (Roley et al. 2014). Greater discharge over the year may drive streams toward heterotrophy by transporting greater amounts of organic matter from riparian soils for microbes to consume (Demars 2019). Additionally, discharge, watershed area, and stream size are often strongly correlated, and others have found GPP and ER to respond positively to drainage area (Mejia et al. 2019). While we did not find a direct relationship between watershed area and discharge in this study, this positive relationship between discharge and ER could potentially be an artifact of stream size. However, there was no relationship between discharge and GPP; thus, if stream size were affecting metabolism, it would be through its influence on heterotrophic and autotrophic carbon processing rather than production. Another possibility may be that grazers are mediating this relationship; herbivores may be more active in streams with lower discharge, thinning algal mats and increasing P:R ratios by ameliorating competition among periphyton, removing senescent cells, and ingesting microbes in the mat (Peckarsky et al. 2015).

The observed relationships between discharge and metabolism appear to be driven more by *Groundwater* than *Runoff* streams. Additionally, *Runoff* streams showed increased respiration then decreased at higher discharges. This could reflect potential flow class-specific differences in algal community composition, in which *Runoff* stream communities respond more variably to increases in discharge, though algal community data and flow-metabolism measurements from a greater number of streams would be needed to support this with certainty.

Greater discharge drove streams to be more heterotrophic due to streamflow stimulating ER with no detectable effect on GPP. This relationship between net ecosystem metabolism (NEM) and discharge may also have resulted from differences in hyporheic connectivity and channel substrate type and heterogeneity, reflecting an important point with respect to flow classifications: even during times of seemingly similar hydrologic conditions, ecosystem structure is an important mediator of functional responses from autotrophs and heterotrophs.

Mean annual discharge was the only flow variable we found to be related to metabolism in this study. This was likely through effects on organic matter transport coupled with the presence or absence of a hyporheic corridor that provided additional space for biota to colonize as well as a source of continuous flow by groundwater intrusion. The importance of hyporheic connectivity and channel substrate to stream metabolism is well-documented, as carbon is utilized from surface water organic matter to support hyporheic respiration (Jones et al. 1995), and while greater discharge could have reduced microbial abundance in *Runoff* streams with no hyporheic refuge, it could have had a stimulatory effect in *Groundwater* streams with subsurface microbes benefitting from organic matter subsidies coming from upstream and the immediate riparian in elevated flows. The hyporheos is a crucial refuge for biota in streams that are susceptible to flash floods as well as drying (Dole-Olivier 2011, Stubbington 2012), and these two flow classifications exhibited distinct disturbance regimes over the course of the study; namely, *Runoff* streams are susceptible to drying, while *Groundwater* streams tend to experience more flood events of greater magnitude. *Groundwater* streams’ connectivity to groundwater may have facilitated greater production and respiration in *Groundwater* streams during certain times of the year by providing asylum for biota to resist to the effects of drying (in the one *Groundwater* stream that dried for nine days) and flooding. In short, discharge and disturbance regimes are different across flow regimes, but hyporheic connectivity may mitigate impacts of drying and flooding on *Groundwater* stream metabolism.

Temperature and light have been shown to synergistically enhance the photosynthetic capacity of primary producers. Temperature can also be predictive of respiration, as it exerts control on the speed of organismal metabolism (Hill et al. 2000, Mulholland et al. 2001, Acuña et al. 2008, Beaulieu et al. 2013). Interestingly, we found the opposite trend-cooler streams had greater rates of respiration. However, this effect of temperature may be related to the volume of groundwater flowing into the system and actually be an indirect measure of groundwater influence (Constantz 1998).

Similar to Bernot et al. (2010), we identified no relationships between phosphorus and GPP. However, other inter-regional studies have determined P to be a driver of GPP, and P concentrations were similar to those reported by others in forested systems (Lamberti and Steinman 1997, Mulholland et al. 2001). Our study sites exhibited low P concentrations; there was not a large gradient in TP concentrations, as these streams were minimally-impacted forested systems. The same was true of nitrogen; total N was also not related to GPP or ER. Others have also found N to be a significant predictor of GPP (Bernot et al. 2010). However, nutrient concentrations were similarly low across sites throughout the study, and our results indicate that the primary influence on metabolism across *Groundwater* and *Runoff* sites was channel discharge.

## CONCLUSION

This comparison of forested systems across flow types provides a foundation for refining comparisons of stream metabolism across systems that may be similar in surrounding land cover, but differ in intermittency, discharge, dominant water source, and hyporheic connectivity. This provides insight into natural variation based on differences in flow versus anthropogenic hydrologic alteration but can also inform predictions regarding how depletion of groundwater resources can influence flow-function relationships. Loss of groundwater inputs will render stream biota more susceptible to droughts and will reduce the capacity to transport organic matter downstream for heterotrophic processing and sustenance of microbial carbon stocks that are important for lotic communities (Battin 1999, Demars 2019).

Our efforts reveal that forested stream metabolism rates are dependent upon discharge and exhibit seasonal trends based on flow regime classification. While biome and land use play key roles in determining stream metabolism, other factors such as streambed stability, hyporheic connectivity, and dominant water source may interact with landscape-level variables to produce variation in carbon dynamics across streams with similar land cover. These controls and flow-function relationships must be integrated across ecosystems as we work to understand sources, fates, and drivers of carbon transformation and transport in aquatic systems (Bernhardt et al. 2017, Hotchkiss et al. 2018).

## ACKNOWLEDGEMENTS

We are grateful to Blake Lefler, Tyler Fletcher, Delaney Hall, Kayla Sayre, Hal Halvorson, Brad Austin, and Richard Walker for field assistance. We warmly thank Brian Haggard, Kristofor Brye, Kusum Naithani for suggestions on earlier versions on the manuscript. This work was funded by the USGS 104b program through the Arkansas Water Resources Center in FY2015. Any use of trade, firm, or product names is for descriptive purposes only and does not imply endorsement by the U.S. Government.

